# MutationalPatterns: The one stop shop for the analysis of mutational processes

**DOI:** 10.1101/2021.11.01.466730

**Authors:** Freek Manders, Arianne M. Brandsma, Jurrian de Kanter, Mark Verheul, Rurika Oka, Markus J. van Roosmalen, Bastiaan van der Roest, Arne van Hoeck, Edwin Cuppen, Ruben van Boxtel

## Abstract

**Background:** The collective of somatic mutations in a genome represents a record of mutational processes that have been operative in a cell. These processes can be investigated by extracting relevant mutational patterns from sequencing data.

**Results:** Here, we present the next version of MutationalPatterns, an R/Bioconductor package, which allows in-depth mutational analysis of catalogues of single and double base substitutions as well as small insertions and deletions. Major features of the package include the possibility to perform regional mutation spectra analyses and the possibility to detect strand asymmetry phenomena, such as lesion segregation. On top of this, the package also contains functions to determine how likely it is that a signature can cause damaging mutations (i.e., mutations that affect protein function). This updated package supports stricter signature refitting on known signatures in order to prevent overfitting. Using simulated mutation matrices containing varied signature contributions, we showed that reliable refitting can be achieved even when only 50 mutations are present per signature. Additionally, we incorporated bootstrapped signature refitting to assess the robustness of the signature analyses. Finally, we applied the package on genome mutation data of cell lines in which we deleted specific DNA repair processes and on large cancer datasets, to show how the package can be used to generate novel biological insights.

**Conclusions:** This novel version of MutationalPatterns allows for more comprehensive analyses and visualization of mutational patterns in order to study the underlying processes. Ultimately, in-depth mutational analyses may contribute to improved biological insights in mechanisms of mutation accumulation as well as aid cancer diagnostics. MutationalPatterns is freely available at http://bioconductor.org/packages/MutationalPatterns.

## Background

Mutational landscapes in the genomes of cells are the result of a balance between mutagenic and DNA-repair processes (1). The somatic mutations that shape these landscapes gradually accumulate throughout life in both healthy and malignant cells (2,3). As a result, the complete collection of somatic mutations in the genome of a cell forms a record of the mutational processes that have been active throughout the life of that cell. In-depth analyses of somatic mutations can allow us to better understand the mutational processes that caused them (4).

First, such analyses can provide insight into the etiology of cancer by identifying mutagenic exposures, which ultimately contribute to the accumulation of cancer driving mutations. For example, we recently identified a mutational pattern caused by a carcinogenic strain of *Escherichia coli* found in the gut of ~20% of healthy individuals (5). This pattern matched mutations found in colorectal cancer driver genes, indicating a direct role in tumorigenesis. Mutational patterns have been systematically determined *in vitro* for many environmental mutagenic agents, which can be used to deduce cancer causes (6). The effects of such agents can also be found *in vivo*. For example, we recently found mutations caused by exposure to the antiviral drug ganciclovir, which patients received to treat a viral infection after a hematopoietic stem cell transplant (7). Second, studying mutational processes can be useful for improved cancer diagnostics. For example, the presence of certain mutational signatures can be used as a functional readout for deficiency of homologous recombination (HR)-mediated double strand break repair (8,9). Cancers with a defect in this repair pathway are selectively sensitive to poly(ADP-ribose) polymerase (PARP) inhibitors, providing a targeted therapy for the patients (10,11).

One of the most popular tools to analyze somatic mutation profiles is the R/Bioconductor package MutationalPatterns, which can be used to easily investigate mutation spectra (12–19). It can also be used to identify new signatures in mutation data using Nonnegative Matrix Factorization (NMF) and to determine the contribution of previously defined signatures to a sample using a method known as “signature refitting” (4). However, the original version of this package has several limitations. First, the package is limited to single base substitutions (SBSs) and cannot be used for small insertions and deletions (indels) or double base substitutions (DBSs) even though signatures for these mutation types have recently been identified in large pan-cancer sequencing efforts (13).

The package also suffers from signature overfitting when determining the contribution of known patterns to a sample, which can result in too many signatures being attributed (20). Additionally, the package only allows for analyzing spectra for mutations in the entire genome, making it difficult to study the involvement of specific genomic elements, such as enhancers or secondary hairpin structures. The ability to investigate the role of such elements in mutation accumulation is important, because this allows for identifying the molecular mechanisms by which certain processes induce mutagenesis (21–23).

Here we present a novel, almost completely rewritten version of MutationalPatterns for the analysis of mutational processes, which is easy-to-use and contains many new features, such as DNA lesion segregation (24). Existing features have also been improved, resulting in a very comprehensive package that can be used for both basic and more advanced mutational pattern analyses. MutationalPatterns supports DBSs, multi base substitutions (MBSs) and indels, and can automatically extract all these mutation types from a single variant call format (VCF) file. The package can generate region specific spectra and signature contributions to study the varying activities of mutational processes across the genome. The package also generates more accurate results by supporting stricter signature refitting. This refitting can also be bootstrapped to determine the confidence of the results. Additionally, a process known as lesion segregation can be investigated.

The MutationalPatterns package can be used to generate novel biological insights, which we demonstrate by applying it to whole genome sequencing (WGS) data obtained from a lymphoblastoid cell line, in which specific DNA repair processes were deleted using CRISPR-Cas9 genome editing, as well as by applying the package on large cancer datasets.

Additionally, we demonstrate that the package scales well on these large datasets. Finally, we show the improved accuracy of the stricter signature refitting using simulated data.

## Implementation

### Mutation profiles

MutationalPatterns supports SBSs, DBSs, MBSs and indels. Multiple mutation types are allowed to be present in a single VCF file so that users do not have to split them beforehand. A specific mutation type can be selected as an argument with the “read_vcfs_as_granges” function when reading in the VCF files. Alternatively, the “get_mut_type” function can be used on data that is already loaded in the memory.

DBS and MBS variants can be called by various variant callers, such as the Genome Analysis Toolkit (GATK) Mutect2, in two different ways (25). The variants can be called explicitly as DBS and MBS variants or as neighboring SBSs. A downside of the first approach is that neighboring germline and somatic mutations can be called as a single combined DBS or MBS, because the variants are compared to the reference instead of the control sample. MutationalPatterns supports both approaches. When the second approach is used, neighboring SBSs will be merged into somatic DBS or MBS variants.

Because they get merged, DBS and MBS variants are no longer incorrectly identified as separate SBSs by MutationalPatterns. This improves the quality of the SBS profiles, as DBS and MBS mutations often have a very different context on account of them being caused by different processes (13) (Additional file 1: Figure S1).

The COSMIC contexts of SBS, indel and DBS variants can be retrieved with fast vectorized functions, namely “mut_context”, “get_indel_context” and “get_dbs_context”. The context of SBS variants consisted of its direct 5’ and 3’ bases in the original package. These contexts were chosen because they are generally the most informative and adding more bases drastically increases the feature space, leading to sparsity (4). Indeed, adding only one extra base to both the upstream and downstream context increases the number of features from 96 to 1536. However, with the increasing availability of large sequencing cohorts such large feature spaces have become more manageable, making it easier to examine nucleotide preference more upstream or downstream of the mutated base. Therefore, MutationalPatterns’ users can now choose any context size for SBSs.

The mutation contexts can be used for custom analyses. Alternatively, the number of mutations per context can be counted, resulting in a count matrix, where each row is a context and each column a sample. These matrices are created with the “mut_matrix”, “mut_matrix_stranded”, “count_indel_contexts”, “count_dbs_contexts” and “count_mbs_contexts” functions. The “count_mbs_contexts” function uses the length of the MBSs, because to date no COSMIC consensus has been defined.

The count matrices can be plotted as spectra or profiles for all the mutation types (Fig. 1a, b, c). The SBS spectra can be displayed in the individual samples. Additionally, the error bars can be displayed as standard deviation, 95% confidence interval (CI) and the standard error of the mean. A count matrix with a larger context can be visualized using the new “plot_profile_heatmap” or “plot_river” functions (Fig. 1d, Additional file 1: Figure S2). This last function can be especially helpful to provide a quick overview of a mutation spectrum with a wider context. Next to visualizing them, a count matrix can also be used for downstream analyses, such as a *de novo* extraction of mutational signatures. In some cases, it can be useful to pool multiple samples within a count matrix to increase statistical power. This can be done using the new “pool_mut_mat” function.

**Fig. 1.**
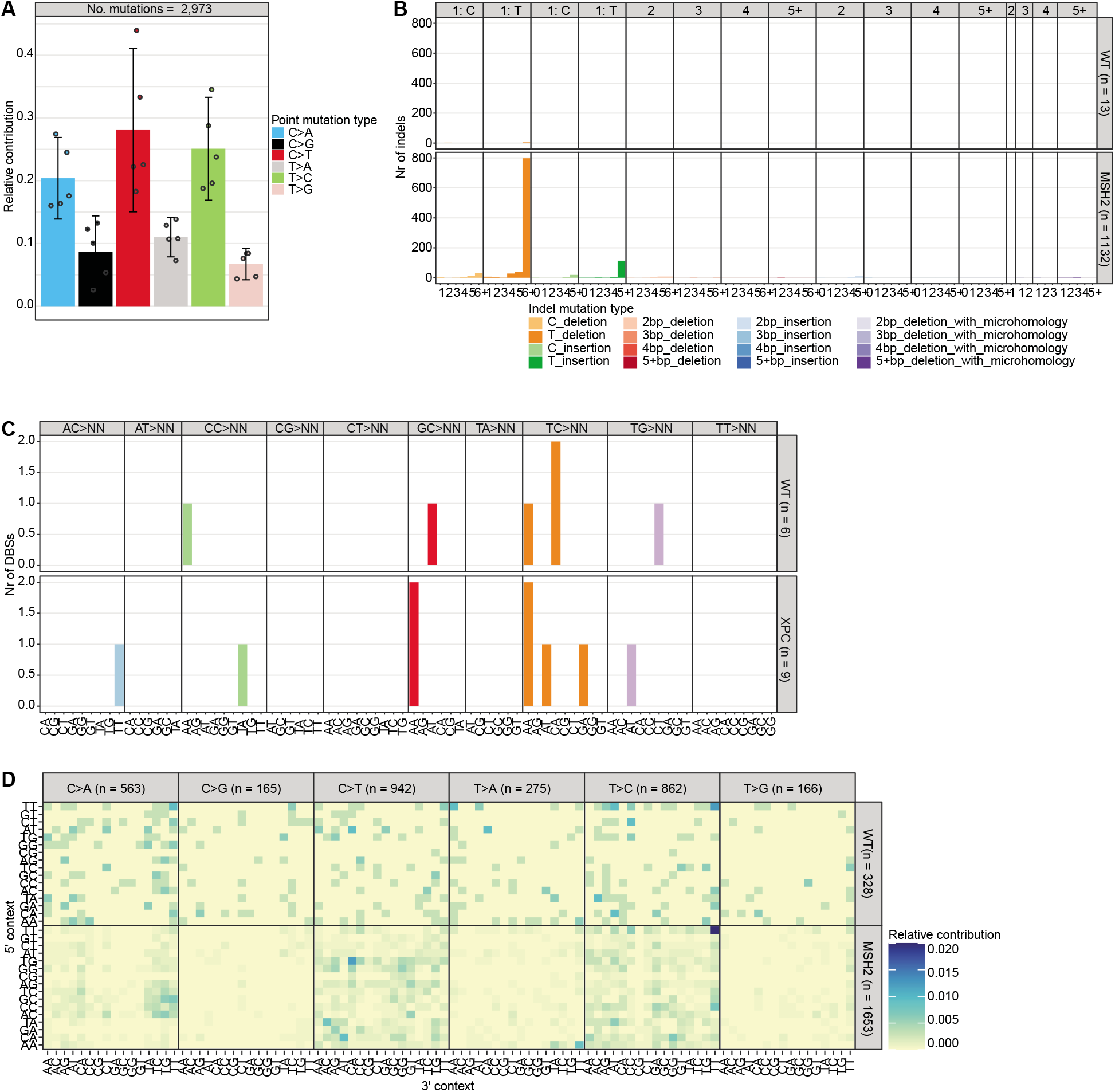
Mutation profiles can be made for multiple mutation types **a** Relative contribution of the indicated mutation types to the point mutation spectrum. Bars depict the mean relative contribution of each mutation type over all the samples and error bars indicate the 95% confidence interval. The dots show the relative contributions of the individual samples. The total number of somatic point mutations per tissue is indicated. **b** Absolute contribution of the indicated mutation types to the indel spectrum for the wild-type (WT) and *MSH2* knockout. The total number of indels per sample is indicated. **c** Absolute contribution of the indicated mutation types to the DBS spectrum for the wild-type (WT) and *XPC* knockout. The total number of DBSs per sample is indicated. **d** Heatmap depicting the relative contribution of the indicated mutation types and the surrounding bases to the point mutation spectrum for the WT and *MSH2* knockout. The total number of somatic point mutations per tissue is indicated.

### Region specific analyses

Mutational processes can be influenced by regional genomic features at multiple scales, such as chromatin landscape, secondary hairpin structures as well as the major and minor groove of the DNA (21–23). With the previous version of MutationalPatterns, it was possible to test for enrichment and/or depletion of the mutation load in such regions. However, the package lacked the possibility to automatically correct for multiple testing. In addition, mutational profiles in genomic regions could not be easily assessed. In MutationalPatterns, multiple testing correction is now automatically performed when testing for enrichment and depletion. In addition, multiple significance levels are now supported, which can be visualized using one or multiple asterisks. Furthermore, regional mutation profiles can be determined in detail. This is done by first splitting mutations based on pre-defined genomic regions, with the new “split_muts_region” function, which requires a GRanges or GRangesList object containing chromosome coordinates as its input. These coordinates can be read into R from file types like “.txt” or “.bed” files or they can be directly read from databases, such as Ensembl (26). This analysis can be performed for multiple samples and multiple types of regions at once. A user could, for example, split a set of mutations into “promoter”, “enhancer” and “other” mutations.

Splitting the mutations according to different genomic regions results in a GRangesList containing sample/region combinations. These combinations can be treated as separate samples by, for example, performing *de novo* signature analysis to identify processes that are specifically active in certain genomic regions. Knowing in which regions a signature is predominantly present, can lead to a better understanding of its etiology. Instead of treating the sample/region combinations as separate samples, the genomic regions can also be incorporated into the mutational contexts, using the new “lengthen_mut_matrix” function. This means that a mutational context like “A[C>A]A” could be split into “A[C>A]A-promoter” and “A[C>A]A-enhancer”. This analysis allows users to generate signatures that contain different mutation contexts in different genomic regions. Such signatures could be more specific than the regular COSMIC signatures.

Region-specific mutation spectra can be visualized with the “plot_spectrum_region” function, which contains the same arguments as the “plot_spectrum” function (Fig. 2a, b). In addition, region-specific 96-channel mutation profiles can be visualized with the new “plot_profile_region” function, which contains the same arguments as the “plot_96_profile” function (Fig. 2c). Both the “plot_spectrum_region” and “plot_profile_region” functions contain a “mode” argument, which allows users to normalize for the occurrence of the different mutation types per sample/region combination, per sample, or not at all.

**Fig. 2.**
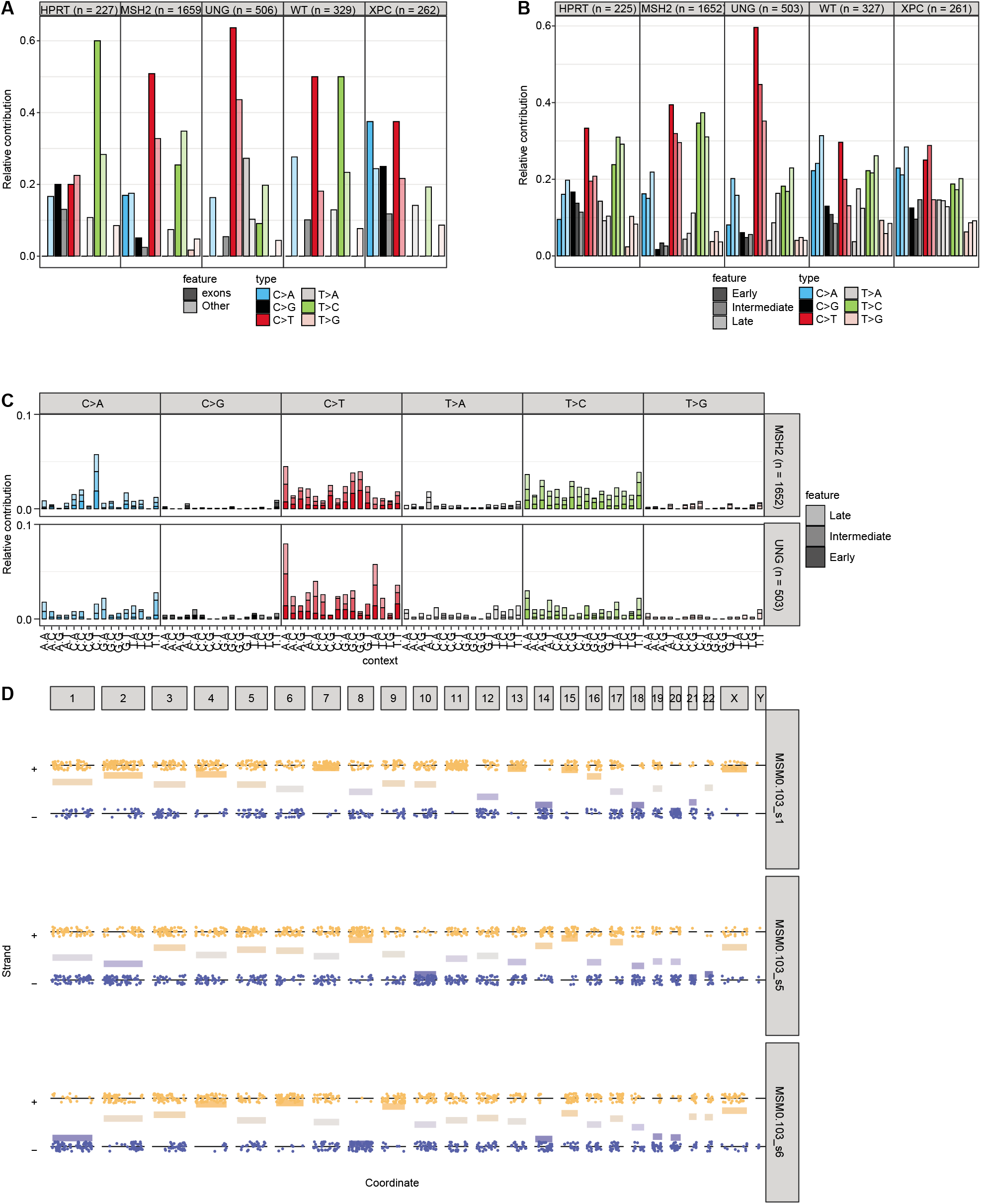
Regional spectra show differences between genomic regions **a** Relative contribution of the indicated mutation types to the point mutation spectrum split between exons and the rest of the genome for each sample. **b** Relative contribution of the indicated mutation types to the point mutation spectrum split between early-, intermediate-, and late-replicating DNA for each sample. **c** Relative contribution of each trinucleotide change to the point mutation spectrum split between early-intermediate and late-replicating DNA for each sample. **d** A jitter plot depicting the presence of lesion segregation for each sample per chromosome. Each dot depicts a single base substitution. Any C>N or T>N is shown as a “+” strand mutation, while G>N and A>N mutations are shown on the “-” strand. The x-axis shows the position of the mutations. The horizontal lines are calculated as the mean of the “+” and strand, where “+” equals 1 and equals 0. They indicate per chromosome on which strand most of the mutations are located. The mutations were downsampled to 33% to reduce the file size.

Instead of using pre-determined genomic regions, it is also possible to compare the mutation spectra of regions with different mutation densities. These regions can be identified using the new “bin_mutation_density” function.

Regional mutational patterns can also be investigated using an unsupervised approach, which is unique to MutationalPatterns, with the new “determine_regional_similarity” function. This function uses a sliding window approach to calculate the cosine similarity between the global mutation profile and the mutation profile of smaller genomic windows, allowing for the unbiased identification of regions with a mutation profile, that differs from the rest of the genome. Users can correct for the oligonucleotide frequency of the genomic windows using the “oligo_correction” argument. The function returns an S4 object, containing the genomic windows with their associated cosine similarities and the settings used to run the function. Because of the unbiased approach of this function, it works best on a large dataset containing at least 100,000 substitutions. The result of this analysis can be visualized using the new “plot_regional_similarity” function.

### Lesion segregation

Mutation spectra sometimes contain Watson versus Crick strand asymmetries (24). These asymmetries can be the result of many DNA lesions occurring during a single cell cycle. If these lesions are not properly repaired before the next genome duplication, then the resulting sister chromatids will segregate into different daughter cells, which will each inherit the lesions on opposite strands. This process is known as lesion segregation (24). The presence of lesion segregation in mutation data can be calculated with the new “calculate_lesion_segregation” function. This calculation can be done for all mutations together or separately for the different mutation contexts. The results can be visualized using the “plot_lesion_segregation” function (Fig. 2d, Additional file 1: Figure S3).

### Mutational signature analysis

When performing signature analyses, it is possible to either extract novel signatures using NMF or to fit previously defined signatures to a mutation count matrix (signature refitting). Both approaches can be applied for all mutation types. By combining count matrices of different types, it is even possible to create a composite signature.

MutationalPatterns now supports a variational Bayesian (Bayes) NMF algorithm from the ccfindR package to help choose the optimal number of signatures, in addition to the regular NMF algorithm (27) (Additional file 1: Figure S4). One challenge with *de novo* signature extraction is that extracted signatures can be very similar to previously defined signatures with known etiology. With the new “rename_nmf_signatures” function, these extracted signatures can be identified using cosine similarity scores and their names can be changed from an arbitrary naming to a custom naming that reflects their similarity to these previously defined signatures.

The original MutationalPatterns package already contained the “fit_to_signatures” function, which finds the optimal combination of signatures to reconstruct a profile and calculates a reconstructed profile based on this combination of signatures. However, this approach could lead to too many signatures being used to explain the data (20). One simple method to reduce this overfitting, which was used in the vignette of the previous version of MutationalPatterns, is to remove all signatures with less than 10 mutations. However, this method, which we will call “regular_10+”, only reduced overfitting slightly. To reduce overfitting, we introduce the new “fit_to_signatures_strict” function. The default backwards selection method of this function iteratively refits a set of signatures to the data, each time removing the signature with the lowest contribution. During each iteration the cosine similarity between the original and reconstructed profile is calculated. The iteration process stops when the change in cosine similarity between two iterations is bigger than the user-specified “max_delta” cutoff (Additional file 1: Figure S5). Users can set the “max_delta” cutoff based on their desired sensitivity and specificity. Stricter refitting, with this method, is comparable to a previously described approach and results in less signatures being chosen when tested on mutation data obtained from cell lines that lack specific DNA repair pathways (Fig. 3a, b; see Additional file 2) (13). The “fit_to_signatures_strict” function also has a best subset selection approach. This method works similarly to the backwards selection approach. However, instead of removing the signature with the lowest contribution, each combination of x signatures is tried. This includes signatures that were not included in a previous iteration. Here, x is the number of signatures used during refitting, which is reduced by one in each iteration step. By default, “fit_to_signatures_strict” uses the backwards selection method, because the best subset method becomes very slow when fitting against more than 10-15 signatures. Therefore, we used the backwards selection method for all “strict” signature refitting analyses in the rest of this manuscript. Another way to reduce overfitting is to only use signatures that are known to be potentially active in your tissue/cells of interest. We recommend using this method in combination with “fit_to_signatures_strict” for optimal results.

**Fig. 3.**
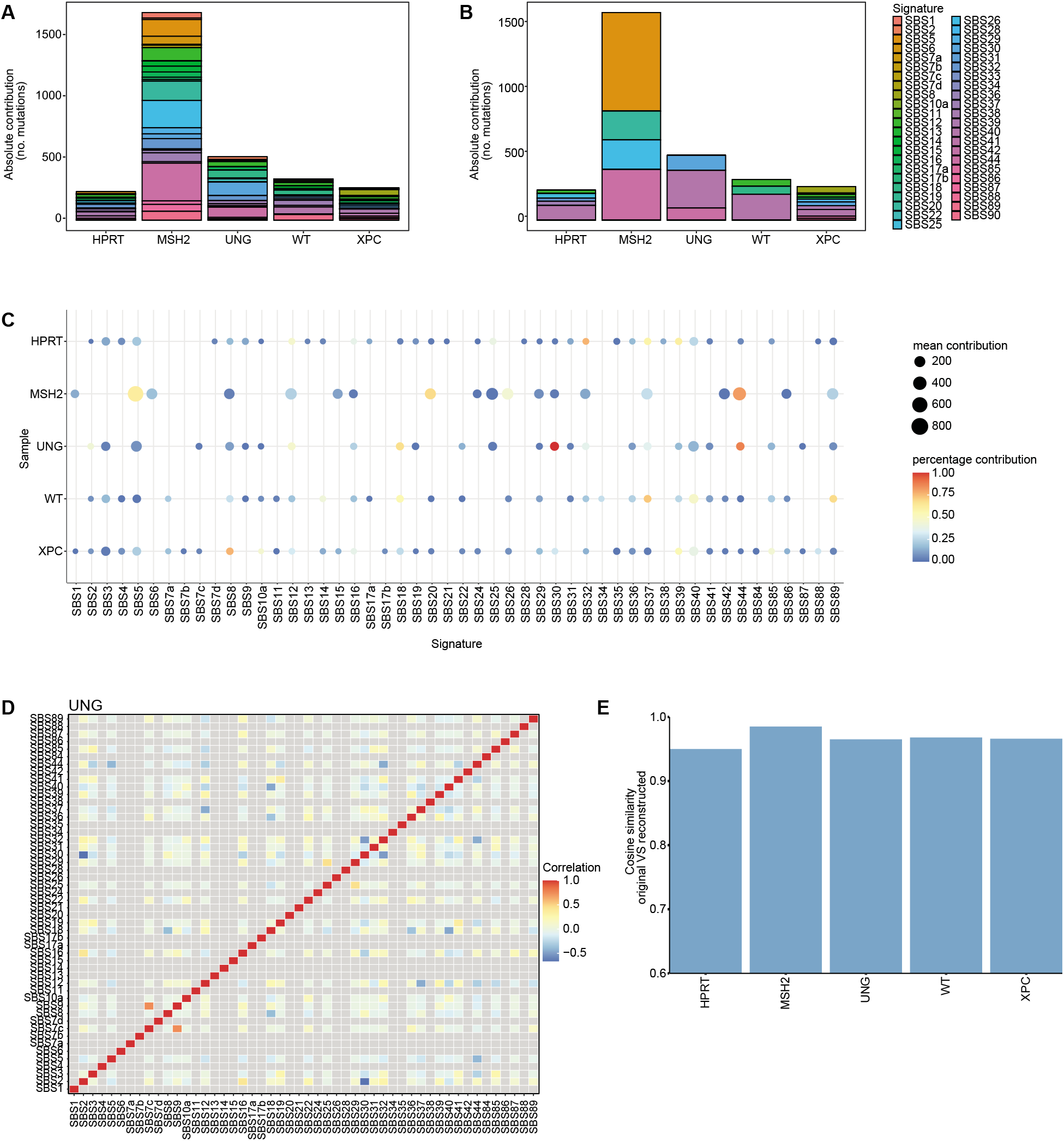
Signature refitting is improved **a** Absolute contribution of each mutational signature for each sample using “regular” signature refitting and **b** “strict” signature refitting. **c** Dot plot showing the contribution of each mutational signature for each sample using bootstrapped signature refitting. The colour of a dot indicates the fraction of bootstrap iterations in which a signature contributed to a sample. The size indicates the mean number of contributing mutations across bootstrap iterations in which the contribution was not zero. **d** Heatmap depicting the Pearson correlation between signature contributions across the bootstrap iterations. **e** Bar graph depicting the cosine similarity between the original and reconstructed profiles of each sample based on signature refitting.

In addition to estimating contributions of signatures to mutation spectra, it is also vital to know how confident these contributions are. The confidence of signature contributions can be determined using a bootstrapping approach with the new “fit_to_signatures_bootstrapped” function, which can use both the strict and the regular refitting methods. Its output can be visualized in multiple ways using the “plot_bootstrapped_contribution” function (Fig. 3c, Additional file 1: Figure S6). The signature contributions can be correlated between signatures across the different bootstrap iterations. This correlation can be visualized using the “plot_correlation_bootstrap” function (Fig. 3d). A negative correlation between two signatures means that each signature had a high contribution in iterations in which the other had a low contribution, which can occur when the refitting process has difficulty distinguishing between two similar signatures. One simple way to deal with highly similar signatures is to merge them. This can be done using the new “merge_signatures” function.

To test the accuracy of signature analysis, the cosine similarity between the reconstructed and original mutation profile needs to be determined. A high cosine similarity between the reconstructed and original profile indicates that the used signatures can explain the original spectrum well. This comparison between reconstructed and original mutation profiles can be visualized with the new “plot_original_vs_reconstructed” function (Fig. 3e).

In order to perform refitting, a matrix is required of the predefined signatures. Signature matrices of the Catalogue of Somatic Mutations in Cancer (COSMIC) (v3.1 + v3.2), SIGNAL (v1) and SparseSignatures (v1) are now included in MutationalPatterns (6,13,15,28). These matrices include general, tissue-specific and drug exposure signatures. The COSMIC matrices also include DBS and indel signatures, next to the standard SBS signatures. Signature matrices can be easily loaded using the new “get_known_signatures” function.

### Signature-specific damaging potential analysis

Some signatures are more likely than others to have functional effects by causing premature stop codons (“stop gain”), splice site mutations or missense mutations, because of sequence specificity underlying these changes. With MutationalPatterns it is now possible to analyze how likely it is for a signature to either cause “stop gain”, “missense”, “synonymous” or “splice site” mutations for a set of genes of interest. For this analysis to be performed, the potential damage first needs to be calculated per mutational context, with the “context_potential_damage_analysis” function. Next, the potential damage per context is combined using a weighted sum to calculate the potential damage per signature using the “signature_potential_damage_analysis” function. The potential damage per signature is also normalized using a “hypothetical” flat signature, which contains the same weight for each mutation context.

This analysis will only take mutational contexts into account. Other features, such as open/closed chromatin, are not considered, because they vary per tissue type. However, this analysis can still give an indication of how damaging a signature might be, which could be supplemented by further custom analyses. This new version of MutationalPatterns also comes with many smaller updates and bugfixes. A comprehensive list can be found in Additional file 3: Table S1.

## Results

### Extended mutation context analysis and regional mutational patterns

To demonstrate the importance of analyzing extended mutation contexts, regional mutational patterns and lesion segregation for characterizing the underlying mutagenic processes, we applied MutationalPatterns to three published mutation datasets. First, we ran MutationalPatterns on 276 melanoma samples from the HMF database. After pooling these samples, we observed that TT[C>T]CT mutations are the most common type of substitution (Fig. 4a). This substitution type is more common than other T[C>T]C substitutions, showing that the extended context has a large effect. Next, we compared the mutation patterns of the melanoma samples between the different genomic regions classified by the Ensembl regulatory build (30). While the patterns look similar, they are significantly different (Fig. 4b) (p = 0.0005, chi-squared test). One subtle difference is the low contribution of T[C>T]A in promoters compared to “Other” regions of the genome, not present in the regulatory build.

**Fig. 4.**
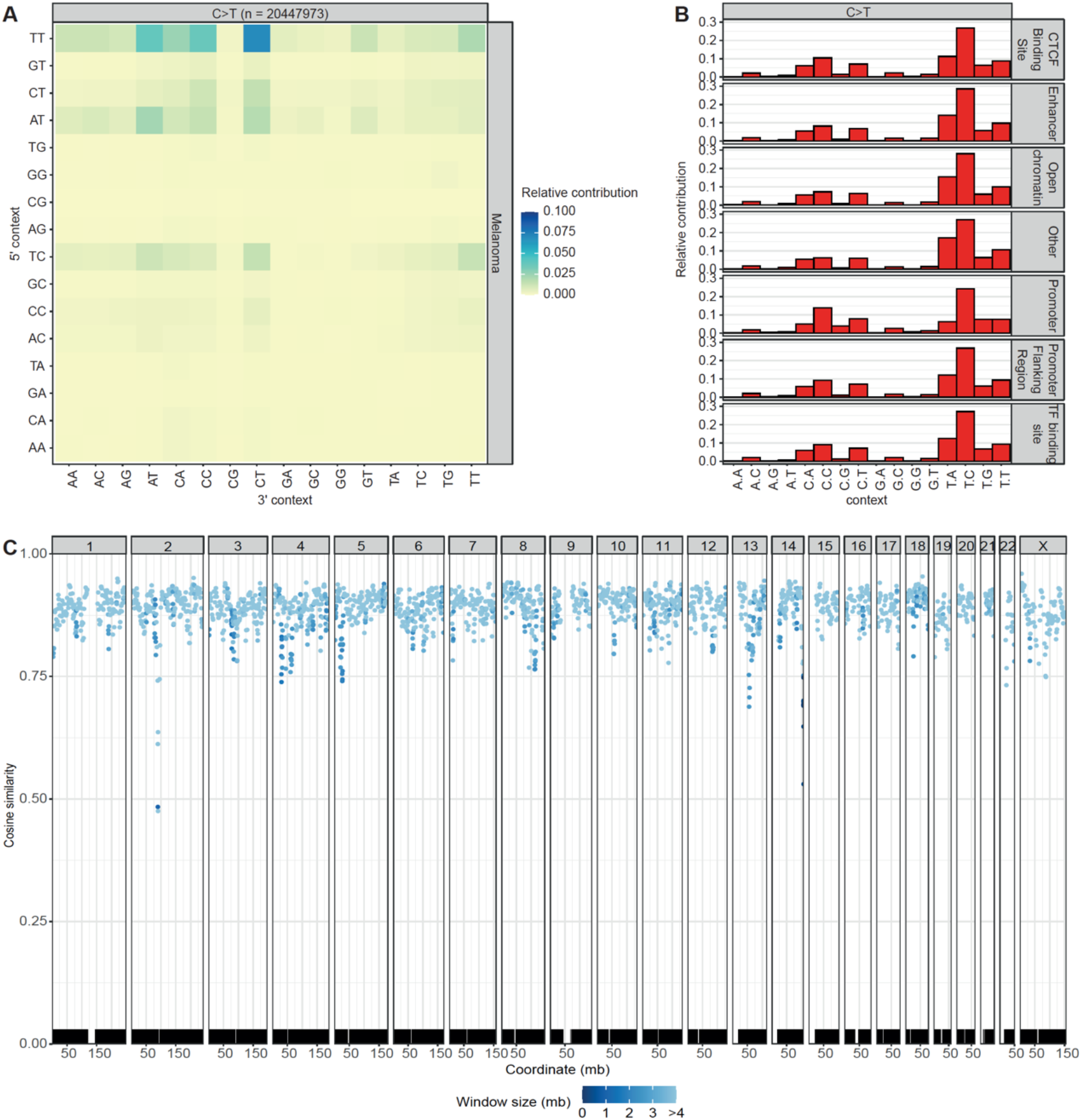
Large cancer datasets show extended and regional mutation patterns **a** Heatmap depicting the relative contribution of the indicated mutation types and the surrounding bases to the point mutation spectrum for metastatic melanomas. The total number of somatic point mutations is indicated. **b** Relative contribution of each C>T trinucleotide change to the point mutation spectrum split between different genomic regions. **c** Graph depicting the similarity in the mutation profile between genomic windows and the rest of the genome. Each dot shows the cosine similarity between the mutation profiles of a single window and the rest of the genome. The dots are colored based on the sizes in mega bases of the windows. The locations of the mutations are plotted on the bottom of the figure.

Next, to show how MutationalPatterns can be used to identify regional activity of specific mutation processes in an unsupervised manner, we applied the package on 217 pooled pediatric B-ALL WGS samples (31). These B-cell-derived leukemias have undergone VDJ recombination, which is associated with somatic hypermutation at loci encoding for immunoglobulin (32,33). As somatic hypermutation is associated with a specific signature, these sites were expected to have a mutation spectrum that is different from the rest of the genome. Indeed, MutationalPatterns was able to detect this for the two VDJ regions, located on chromosomes 2 and 14 (Fig. 4c). Some other regions also seem to have a different mutational pattern, several of which contain PCDH genes. However, further research is needed to explain these results. This example shows how MutationalPatterns can identify region-specific mutational processes in an unsupervised manner.

Finally, to show how MutationalPatterns can identify lesion segregation, we applied it on a dataset known to contain this phenomenon. We found significant lesion segregation in data obtained from induced pluripotent stem cells treated with 0.109 uM of dibenz[a,h]anthracene diol-epoxide (6,24), using the “plot_lesion_segregation” function of MutationalPatterns (Fig. 2d). It was even possible to spot sister-chromatid-exchange events, such as on chromosome 2 of sample MSM0.103_s6 (Fig. 2d, lower panel). To reduce the file size of the figure, 66% of the mutations of each sample were removed using the “downsample” argument of this function. Using MutationalPatterns, we also found lesion segregation in patients that received the antiviral drug ganciclovir (7).

### MutationalPatterns offers more functionality than other mutation analysis tools

An overview of the functions of MutationalPatterns and related tools is shown in Table 1. The original version of MutationalPatterns is also included in this table. An important advantage of the original package was that it combined many mutational analyses into a single package. This new version improves many of these features and adds many new and unique features.

### Mutation matrices can be generated faster

To make MutationalPatterns scalable to large cancer datasets and suitable for interactive analysis we improved the runtime of the “mut_matrix” and “mut_matrix_stranded” functions by vectorizing them. The new functions for retrieving the mutation contexts and generating the mutation matrices have also been written in a vectorized way. As a result, these functions have O(n) or better scaling as tested on a large WGS database from the Hartwig Medical Foundation (HMF) (Additional file 1: Figure S7) (29).

To test their improved performance, we benchmarked the “mut_matrix” and “mut_matrix_stranded” functions on the example data provided in the previous version of MutationalPatterns (Additional file 1: Figure S8). These functions are now respectively 3.4 and 2.6 times as fast on average. In other words, a mutation matrix for 1 million SBSs can now be made in only 135 seconds on a laptop, which makes these functions suitable for large cancer datasets.

**Table 1:**
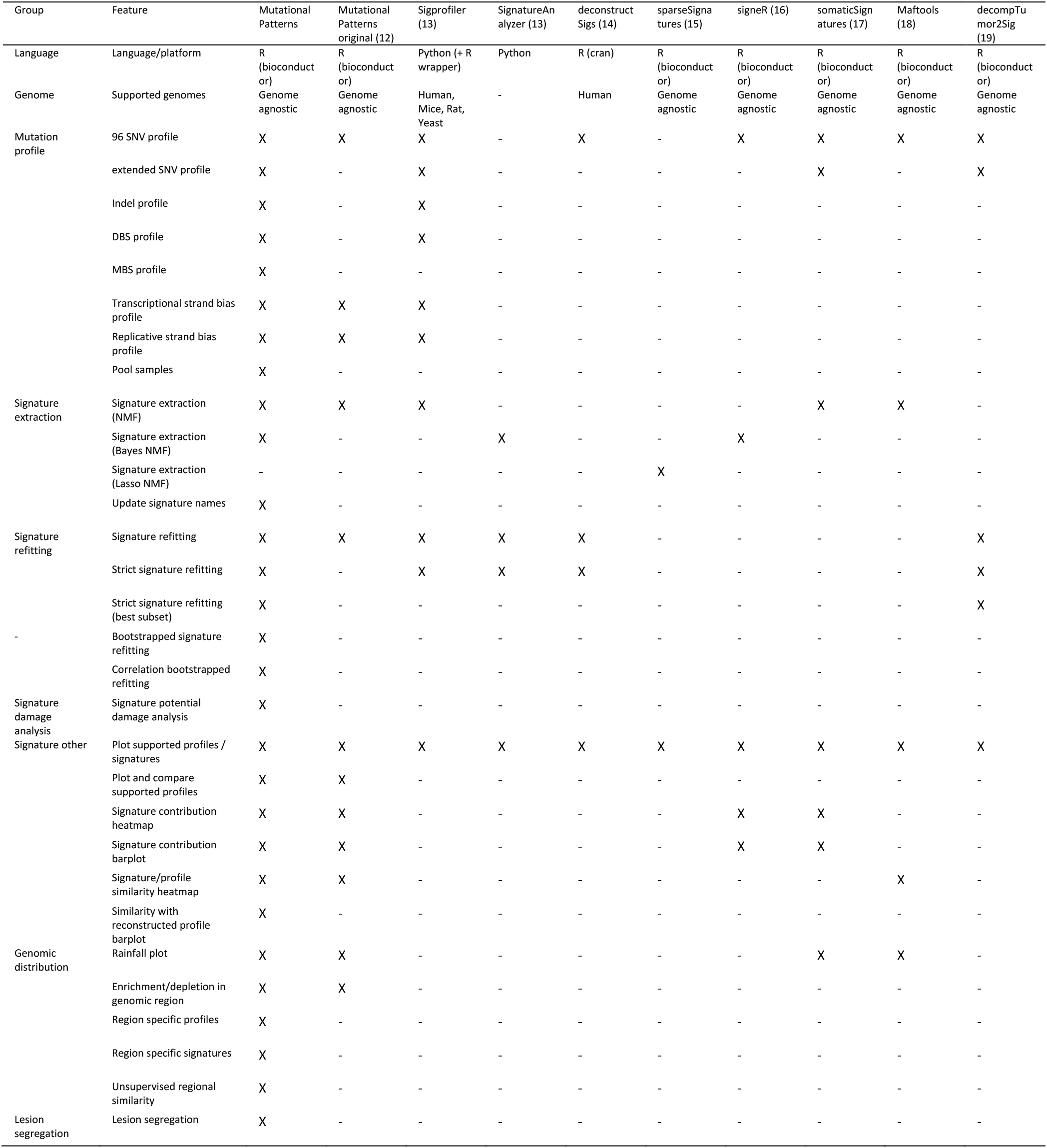
Feature comparison with other packages

### Strict signature refitting improves performance

To determine how well the strict refitting method of MutationalPatterns performs as compared to the regular method, we used simulated mutation matrices. These matrices were generated by sampling trinucleotide changes of 4 different randomly selected signatures. This process was repeated 300 times per matrix, to generate 300 “samples”. Each of the samples in a matrix contained the same number of mutations per signature but was composed of different signatures. The signatures were selected from the first 30 signatures of the COSMIC signature matrix. We limited our analysis to the first 30, because these are the signatures that are most often observed in cancers and therefore more accurately resemble real-life scenarios. In addition, this approach better resembles how the package is used, because users will often fit against a limited number of signatures associated with a specific tissue. By limiting ourselves to the first 30 COSMIC signatures we also reduced overfitting. Any overfitting we observed was thus not caused by us using an unusually large signature matrix. In total we generated 4 matrices, each containing 300 samples. The number of mutations per sample was respectively 200, 400, 2000 and 4000 for the 4 different matrices.

The fraction of correctly attributed mutations to the specific signatures was increased with the strict refitting approach of MutationalPatterns as compared to “regular” or “regular_10+” refitting (Additional file 1: Figure S9a). All the tested refitting methods work better when there are more mutations per signature. Instead of using the number of correctly attributed mutations as a readout for performance, we determined whether the presence and absence of specific signatures was correctly classified. This readout might be more informative for mutational signature analysis because the presence of a signature can be a clinically relevant finding. The strict refitting method achieved a much higher precision than the original methods, while retaining a high correct recall rate (sensitivity) (Additional file 1: Figure S9b). The strict method obtained an area under the curve (AUC) of 0.925, even when only 50 mutations were present per signature, indicating that refitting can be performed on relatively small amounts of mutations.

### SBS10a and SBS18 have a high damage potential

We applied the “signature_potential_damage_analysis” function on the COSMIC signatures. This analysis showed that SBS10a and SBS18 are respectively 3.6 and 2.0 times as likely to cause a “stop gain” mutation compared to a completely flat signature, containing the same weight for each mutation context, on a set of genes associated with cancer (Additional file 3: Table S2, Table S3). SBS18 is related to oxidative stress, suggesting that this type of stress has a high potency of generating premature stop codons in genes that are recurrently associated with tumorigenesis (13). In contrast, the clock-like signature SBS1, which also occurs in healthy cells, was 0.81 and 0.40 times as likely to cause “stop gain” and “splice site” mutations, respectively, as compared to a completely flat hypothetical signature (2,34) (Additional file 3: Table S2). The damaging potential of this ageing-related mutational process is thus relatively low. Overall, C>A heavy signatures, like the recently identified ganciclovir signature, have more damage potential, because they are most likely to introduce a premature stop codon in an open reading frame (7). Being able to quickly assess the damage potential of existing and novel signatures can be very useful to prioritize samples and mutagenic exposures for further investigation.

### Applying MutationalPatterns on mutation data of DNA repair-deficiencies

To illustrate the functionality of MutationalPatterns on real-life data and to obtain novel biological insights, we applied it to mutation data obtained from cell lines in which we deleted specific DNA repair pathways using CRISPR-Cas9 genome editing technology (Additional file 1: Figure S10, Additional file 2). In AHH-1 cells, a lymphoblastoid cell line, we generated bi-allelic knockout lines of *MSH2*, *UNG* and *XPC* by transfecting the cells with a plasmid containing Cas9 and a single gRNA against the gene of interest. By co-transfection with a *HPRT-targeting* plasmid, we were able to select the transfected cells using 6-thioguanine, to which only HPRT-sufficient cells are sensitive. Using this protocol, no targeting vectors for each gene of interest were required. We analyzed somatic mutations in *HPRT-only* knockout lines as well as the combination of *HPRT* with *MSH2*, *UNG* and *XPC* (Additional file 2). To catalogue mutations that were acquired specifically in the absence of the targeted DNA repair gene, we used a previously developed method (35). In brief, whole genome sequencing was performed on generated clones and subclones. By subtracting variants present in the clones from those in the subclones, the somatic mutations, that accumulated in between the clonal steps, were determined.

The SBS profiles are shown in Additional file 1: Figure S11. Interestingly, the profile observed in the *MSH2* knockout cell line displayed a large C[C>A]T peak. When extending the sequence context surrounding the mutated base, the *MSH2* deficiency profile showed a large TT[T>C]TT peak, suggesting that this extended context surrounding mutated thymine residues is important for the underlying mutagenic process (Fig. 1d).

Next, we examined regional mutation patterns. The spectra of the *MSH2*- and *UNG*-deficient cells varied between the exonic regions and the rest of the genome (Fig. 2a)(fdr = 0.0012, fdr = 0.0012; chi-squared test). Their exons contained more C>T and less T>C mutations. The other samples did not show a significant difference in regional mutation spectra. However, when we downsampled all the samples to 227 mutations, which is the number of mutations in the *HPRT* only knockout, no significant regional mutation patterns were observed in *MSH2* and *UNG* knockout cells. This suggests that with this number of mutations insufficient statistical power was obtained for these analyses. Next to examining mutation profiles in exonic regions, we also analyzed regions with different replication timing dynamics, using the median replication timing data from 5 B-lymphocyte cell lines from ENCODE (Fig. 2b, Additional file 3: Table S4) (40). The spectra of *MSH2* and *UNG* knockouts were different between early-, intermediate- and late-replicating DNA (fdr = 0.0012, fdr = 0.0012; chi-squared test). Early replicating DNA has more C>T and less C>A than late replicating DNA. These differences were still present when downsampling was applied (fdr = 0.0025, fdr = 0.010; chi-squared test). Based on these region-specific analyses, we can conclude that the mutational processes active in the *MSH2* and *UNG* knockouts show varying activities in different regions of the genome, a result that cannot easily be obtained with other tools.

We also tested if any of the DNA repair knockout cells displayed lesion segregation, which would indicate that most of the mutations occurred during a single cell-cycle; however, this was not the case (Additional file 1: Figure S6).

Finally, we looked at the mutational signatures in the knockout samples. Based on signature refitting, the *MSH2* knockout contained contributions of SBS5, SBS20, SBS26 and SBS44 (Fig. 3b, c). Because of the bootstrapping we can be more confident in these results. SBS5 is a clock-like signature, with unknown etiology. SBS20, SBS26 and SBS44 are all associated with defective DNA mismatch repair in cancer mutation data (13). The UNG knockout contained contributions from SBS30, which has previously been attributed to deficiency of the base excision repair gene *NTHL1* (13). The glycosylase encoded by *NTHL1* is involved in the removal of oxidized pyrimidines from the DNA and therefore SBS30 likely reflects an alternative consequence of oxidative stress-induced mutagenesis as compared to SBS18. However, *UNG* is a glycosylase that is believed to remove uracil residues from the DNA (36,37). Therefore, our data suggests that SBS30 can be caused, besides oxidized pyrimidines, by unremoved uracil residues. Alternatively, *UNG* may also, to a certain extent, be involved in the removal of oxidized pyrimidines from the DNA. Even though the contribution of SBS30 was relatively modest in the *UNG* knockout, it was consistently picked up by the bootstrapping algorithm. This observation indicated that the number of mutations attributed to a signature is not necessarily related to the confidence of its presence, which further demonstrates the importance of our bootstrapping approach. Unexpectedly, the contribution of SBS30 in *UNG* knockout cells was negatively correlated with SBS2, even though their cosine similarity is only 0.46 (Fig. 3d). This indicates that the refitting algorithm has difficulty choosing between SBS2 and SBS30. Such difficulties in signature selection could lead to different and possibly incorrect signatures being attributed to similar sample types. Understanding the correlation of estimated signature contributions between different signatures, which can be achieved with bootstrapping, is important to prevent incorrect interpretation of the data. The *XPC* knockout contained contributions from SBS8. The etiology of this signature is not yet known. However, this finding further confirms the association of SBS8 with nucleotide excision repair deficiency (38,39). Overall, the COSMIC signatures could explain the mutation profiles of most samples quite well, even when strict refitting was used (Fig. 3e).

Next, we studied the indel signatures in these knockout lines. Deletion of *MSH2* resulted in an increased number of indels as compared to wild-type cells (Fig. 1b). Most of these indels were single thymine deletions in thymine mononucleotide repeat regions. Signature analysis indicated that ID1, ID2 and ID7 contributed to the indel pattern in the MSH2-deficient cells (Fig. 5a, b). Of these, ID1 and ID2 are associated with polymerase slippage during DNA replication and found in large numbers in cancers with mismatch repair deficiency. ID7 is also associated with defective DNA mismatch repair, but not attributed to polymerase slippage (13). Together these signatures could explain the mutational indel profile of *MSH2* knockout cells very well (Fig. 5c), showing that MutationalPatterns can perform indel signature refitting. None of the knockout cells displayed a strongly increased number of DBSs as compared to the wild-type cells (Fig. 1c).

**Fig. 5.**
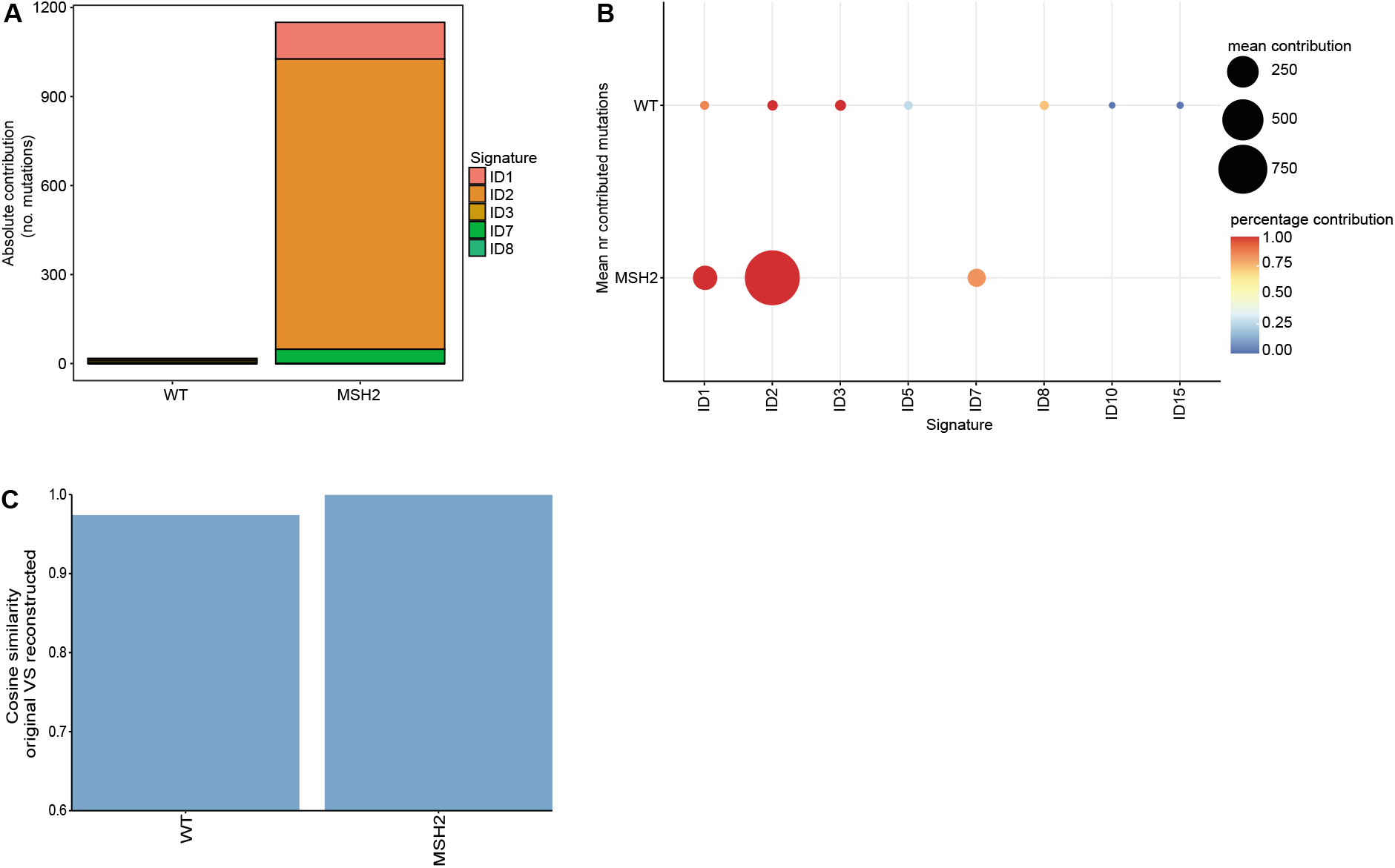
Indel signatures can explain the MSH2 profile **a** Relative contribution of each mutational signature for the wild-type (WT) and *MSH2* samples using strict signature refitting. **b** Dot plot showing the contribution of each mutational signature for the WT and *MSH2* samples using bootstrapped signature refitting. The color of a dot indicates the fraction of bootstrap iterations in which a signature contributed to a sample. The size indicates the mean number of contributing mutations across bootstrap iterations in which the contribution was not zero. **c** Bar graph depicting the cosine similarity between the original and reconstructed profiles of the WT and *MSH2* samples based on signature refitting.

## Discussion

The novel version of MutationalPatterns has been designed to be easy-to-use in such a way that both experienced bioinformaticians and wet-lab scientists with a limited computational background can use it. The code is written in the tidyverse style, which makes it more similar to natural English and therefore easier to understand for non-programmers. MutationalPatterns gives clear error messages with tips on how to solve them, in contrast to the default error messages in R, which can sometimes be cryptic. The updated vignette, accompanying the package, not only explains how the functions in the package can be used, but also informs users on the pros and cons of the different analysis strategies.

Similar to the previous version of the package, plots are all generated using ggplot2 (41). This allows users to visualize their data in highly customizable plots that can be easily modified. Because this feature was not readily apparent for many users of the original MutationalPatterns package, we have now explicitly showed how to modify the elements of a plot, such as the axis and theme, in the vignette.

We have adopted unit testing for this version of the package, resulting in more than 90% code coverage. This will improve the stability of the package and makes it easier to maintain.

The novel version of MutationalPatterns is already available on Bioconductor as an update of the previous version. MutationalPatterns does not break existing scripts and pipelines, because backwards incompatible changes have been kept to a minimum.

## Conclusions

MutationalPatterns is an easy-to-use R/Bioconductor package that allows in-depth analysis of a broad range of patterns in somatic mutation catalogues, supporting single and double base substitutions as well as small insertions and deletions. Here, we have described the new and improved features of the package and shown how the package performs on existing cancer data sets and on mutation data obtained from cell lines in which specific DNA repair genes are deleted. These analyses demonstrate how the package can be used to generate novel biological insights.

Mutational pattern analyses have proven to be a powerful approach to dissect mutational processes that have operated in cancer and to support treatment decision making in personalized medicine. Therefore, mutational patterns hold a great promise for improved future cancer diagnosis. The MutationalPatterns package can be used to fulfill this promise and we are confident that it will be embraced by the community.

## Availability and requirements

The availability and requirements are listed as follows:

Project name: MutationalPatterns

Project home page: https://github.com/ToolsVanBox/MutationalPatterns

Archived version: https://bioconductor.org/packages/3.14/bioc/html/MutationalPatterns.html

Operating system(s): Linux, Windows or MacOS

Programming language: R (version > = 4.1.0)

License: MIT

## List of abbreviations

HR: homologous recombination
Indels: Insertions and deletions
DBS: double base substitutions
VCF: variant call format
MBS: Multi base substitutions
COSMIC: Catalogue of Somatic Mutations in Cancer
NMF: non-negative matrix factorization
Bayes: Bayesian
AUC: Area under the curve
PCA: Principal component analysis
CI: Confidence interval
WT: wild-type
Mb: mega bases

## Declarations

### Ethics approval and consent to participate

Not applicable

### Consent for publication

Not applicable

### Availability of data and materials

The datasets supporting this article are available on EGA under accession number (Study ID EGAS00001004789).

Additionally, the VCF files and scripts that can be used to reproduce all figures in this paper can be found at https://github.com/ToolsVanBox/MutationalPatterns_manuscript2_data_scripts/

## Authors’ contributions

F.M., R.v.B. and A.M.B wrote the manuscript. F.M. and J.d.K. developed and implemented the package. F.M. and R.O. maintain the package. A.M.B. and M.V. generated the data. F.M. and M.J.v.R. analyzed the data. A.v.H., B.v.d.R. and E.C. tested the package and provided feedback. All authors read and approved the final manuscript.

## Acknowledgements

We would like to thank Francis Blokzijl and Roel Janssen for developing and maintaining the first version of this package. We also want to thank Roel Janssen for his support during the handover of the package. Finally, we would like to thank anybody who tested the package for their feedback.

---

Additional file 1:

PDF (.pdf)

Additional figures

A PDF file containing the additional figures.

Additional file 2:

PDF (.pdf)

Additional methods

A PDF file describing the generation and sequencing analysis of the knockout lines.

Additional file 3:

Excel (.xlsx)

Additional tables

An Excel file containing the additional tables.

## References

1. Helleday T, Eshtad S, Nik-Zainal S. Mechanisms underlying mutational signatures in human cancers. Nat Rev Genet. 2014;15:585–98.

2. Blokzijl F, de Ligt J, Jager M, Sasselli V, Roerink S, Sasaki N, et al. Tissue-specific mutation accumulation in human adult stem cells during life. Nature. 2016;538:260–4.

3. Campbell PJ, Getz G, Korbel JO, Stuart JM, Jennings JL, Stein LD, et al. Pan-cancer analysis of whole genomes. Nature [Internet]. 2020;578(7793):82–93. Available from: https://doi.org/10.1038/s41586-020-1969-6

4. Alexandrov LB, Nik-Zainal S, Wedge DC, Campbell PJ, Stratton MR. Deciphering Signatures of Mutational Processes Operative in Human Cancer. Cell Rep. 2013;3:246–59.

5. Pleguezuelos-Manzano C, Puschhof J, Rosendahl Huber A, van Hoeck A, Wood HM, Nomburg J, et al. Mutational signature in colorectal cancer caused by genotoxic pks+ E. coli. Nature. 2020;580:269–73.

6. Kucab JE, Zou X, Morganella S, Joel M, Nanda AS, Nagy E, et al. A Compendium of Mutational Signatures of Environmental Agents. Cell. 2019;177:821–836.e16.

7. de Kanter JK, Peci F, Bertrums E, Rosendahl Huber A, van Leeuwen A, van Roosmalen MJ, et al. Antiviral treatment causes a unique mutational signature in cancers of transplantation recipients. Cell Stem Cell [Internet]. 2021; Available from: https://www.sciencedirect.com/science/article/pii/S1934590921003374

8. Davies H, Glodzik D, Morganella S, Yates LR, Staaf J, Zou X, et al. HRDetect is a predictor of BRCA1 and BRCA2 deficiency based on mutational signatures. Nat Med. 2017;23:517–25.

9. Nguyen L, W. M. Martens J, Van Hoeck A, Cuppen E. Pan-cancer landscape of homologous recombination deficiency. Nat Commun. 2020;11:5584.

10. Bryant HE, Schultz N, Thomas HD, Parker KM, Flower D, Lopez E, et al. Specific killing of BRCA2-deficient tumours with inhibitors of poly(ADP-ribose) polymerase. Nature. 2005;434:913–7.

11. Fong PC, Boss DS, Yap TA, Tutt A, Wu P, Mergui-Roelvink M, et al. Inhibition of Poly(ADP-Ribose) Polymerase in Tumors from BRCA Mutation Carriers. N Engl J Med. 2009;361:557–68.

12. Blokzijl F, Janssen R, van Boxtel R, Cuppen E. MutationalPatterns: comprehensive genome-wide analysis of mutational processes. Genome Med. 2018;

13. Alexandrov LB, Kim J, Haradhvala NJ, Huang MN, Tian Ng AW, Wu Y, et al. The repertoire of mutational signatures in human cancer. Nature. 2020;578:94–101.

14. Rosenthal R, McGranahan N, Herrero J, Taylor BS, Swanton C. deconstructSigs: delineating mutational processes in single tumors distinguishes DNA repair deficiencies and patterns of carcinoma evolution. Genome Biol. 2016;17:31.

15. Ramazzotti D, Lal A, Liu K, Tibshirani R, Sidow A. De Novo Mutational Signature Discovery in Tumor Genomes using SparseSignatures. bioRxiv. 2019;384834.

16. Rosales RA, Drummond RD, Valieris R, Dias-Neto E, Da Silva IT. signeR: An empirical Bayesian approach to mutational signature discovery. Bioinformatics. 2017;33:8–16.

17. Gehring JS, Fischer B, Lawrence M, Huber W. SomaticSignatures: Inferring mutational signatures from single-nucleotide variants. Bioinformatics. 2015;31:3673–5.

18. Mayakonda A, Lin D-C, Assenov Y, Plass C, Koeffler HP. Maftools: efficient and comprehensive analysis of somatic variants in cancer. Genome Res. 2018/10/19. 2018 Nov;28:1747–56.

19. Krüger S, Piro RM. decompTumor2Sig: identification of mutational signatures active in individual tumors. BMC Bioinformatics [Internet]. 2019;20(4):152. Available from: https://doi.org/10.1186/s12859-019-2688-6

20. Maura F, Degasperi A, Nadeu F, Leongamornlert D, Davies H, Moore L, et al. A practical guide for mutational signature analysis in hematological malignancies. Nat Commun. 2019;10:2969.

21. Polak P, Karlic R, Koren A, Thurman R, Sandstrom R, Lawrence MS, et al. Cell-of-origin chromatin organization shapes the mutational landscape of cancer. Nature. 2015;518:360–4.

22. Buisson R, Langenbucher A, Bowen D, Kwan EE, Benes CH, Zou L, et al. Passenger hotspot mutations in cancer driven by APOBEC3A and mesoscale genomic features. Science (80-). 2019;364:eaaw2872.

23. Gonzalez-Perez A, Sabarinathan R, Lopez-Bigas N. Local Determinants of the Mutational Landscape of the Human Genome. Cell. 2019;177:101–14.

24. Aitken SJ, Anderson CJ, Connor F, Pich O, Sundaram V, Feig C, et al. Pervasive lesion segregation shapes cancer genome evolution. Nature. 2020;583:265–70.

25. Benjamin D, Sato T, Cibulskis K, Getz G, Stewart C, Lichtenstein L. Calling Somatic SNVs and Indels with Mutect2. bioRxiv. 2019;861054.

26. Yates AD, Achuthan P, Akanni W, Allen J, Allen J, Alvarez-Jarreta J, et al. Ensembl 2020. Nucleic Acids Res. 2020;48:D682–8.

27. Woo J, Winterhoff BJ, Starr TK, Aliferis C, Wang J. De novo prediction of cell-type complexity in single-cell RNA-seq and tumor microenvironments. Life Sci Alliance. 2019;2:e201900443.

28. Degasperi A, Amarante TD, Czarnecki J, Shooter S, Zou X, Glodzik D, et al. A practical framework and online tool for mutational signature analyses show inter-tissue variation and driver dependencies. Nat cancer. 2020;1:249–63.

29. Priestley P, Baber J, Lolkema MP, Steeghs N, de Bruijn E, Shale C, et al. Pan-cancer whole-genome analyses of metastatic solid tumours. Nature. 2019;575:210–6.

30. Zerbino DR, Wilder SP, Johnson N, Juettemann T, Flicek PR. The ensembl regulatory build. Genome Biol [Internet]. 2015 Mar;16:56. Available from: http://europepmc.org/articles/PMC4407537

31. Ma X, Liu Y, Liu Y, Alexandrov LB, Edmonson MN, Gawad C, et al. Pan-cancer genome and transcriptome analyses of 1,699 paediatric leukaemias and solid tumours. Nature. 2018 Feb;

32. Chi X, Li Y, Qiu X. V(D)J recombination, somatic hypermutation and class switch recombination of immunoglobulins: mechanism and regulation. Immunology [Internet]. 2020/02/27. 2020 Jul;160(3):233–47. Available from: https://pubmed.ncbi.nlm.nih.gov/32031242

33. Di Noia JM, Neuberger MS. Molecular Mechanisms of Antibody Somatic Hypermutation. Annu Rev Biochem [Internet]. 2007 Jun 7;76(1):1–22. Available from: https://doi.org/10.1146/annurev.biochem.76.061705.090740

34. Alexandrov LB, Jones PH, Wedge DC, Sale JE, Campbell PJ, Nik-Zainal S, et al. Clock-like mutational processes in human somatic cells. Nat Genet. 2015;47:1402–7.

35. Drost J, van Boxtel R, Blokzijl F, Mizutani T, Sasaki N, Sasselli V, et al. Use of CRISPR-modified human stem cell organoids to study the origin of mutational signatures in cancer. Science (80-). 2017;238:eaao3130.

36. Prasad A, Wallace SS, Pederson DS. Initiation of Base Excision Repair of Oxidative Lesions in Nucleosomes by the Human, Bifunctional DNA Glycosylase NTH1. Mol Cell Biol. 2007;27:8442 LP–8453.

37. Li J, Braganza A, Sobol RW. Base Excision Repair Facilitates a Functional Relationship Between Guanine Oxidation and Histone Demethylation. Antioxid Redox Signal. 2013;18:2429–43.

38. Jager M, Blokzijl F, Kuijk E, Bertl J, Vougioukalaki M, Janssen R, et al. Deficiency of nucleotide excision repair is associated with mutational signature observed in cancer. Genome Res. 2019;29:1067–77.

39. Yurchenko AA, Padioleau I, Matkarimov BT, Soulier J, Sarasin A, Nikolaev S. XPC deficiency increases risk of hematologic malignancies through mutator phenotype and characteristic mutational signature. Nat Commun [Internet]. 2020;11(1):5834. Available from: https://doi.org/10.1038/s41467-020-19633-9

40. Moore JE, Purcaro MJ, Pratt HE, Epstein CB, Shoresh N, Adrian J, et al. Expanded encyclopaedias of DNA elements in the human and mouse genomes. Nature. 2020;583:699–710.

41. Wickham H. ggplot2: Elegant Graphics for Data Analysis. Springer-Verlag New York; 2016.

