## Additional file 1 for "MutationalPatterns: The one stop shop for the analysis of mutational processes"

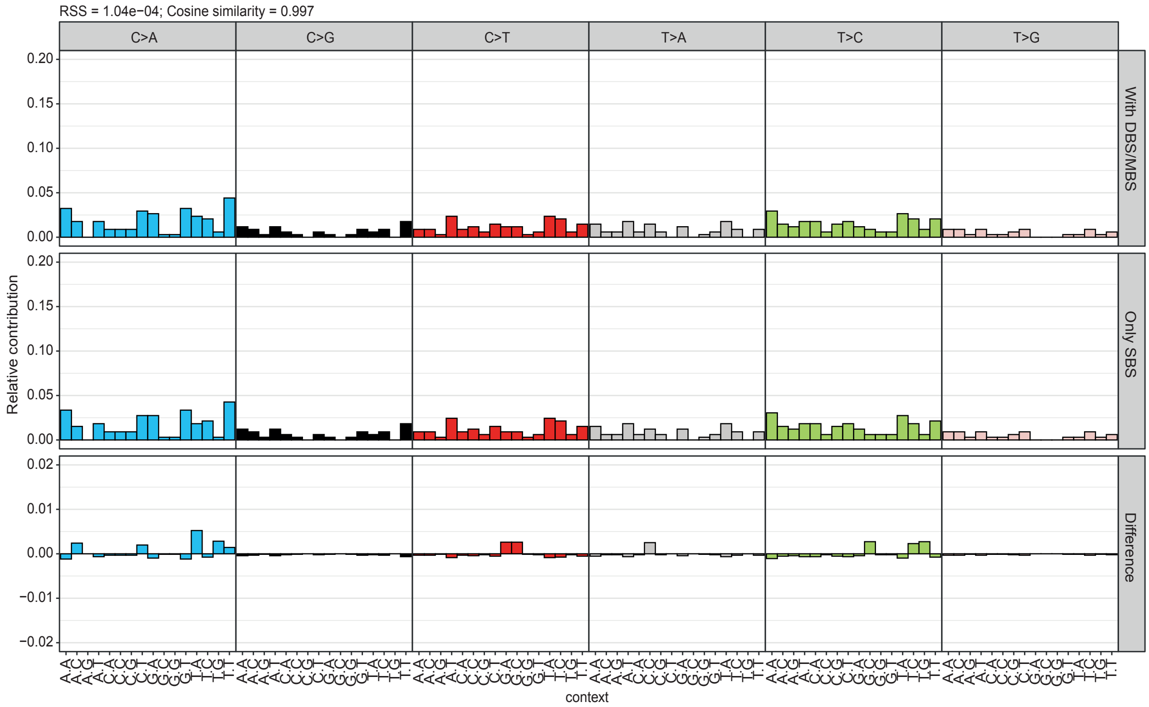


Fig. S1 Removing the DBSs and MBSs results in an improved SBS profile

Relative contribution of each of the 96 trinucleotide changes to the mutational profiles of the wild-type sample. The upper panel shows the profile when DBSs and MBSs are incorrectly classified as SBSs. The middle panel shows the profile with only the SBSs. The lower panel shows the difference between these profiles.

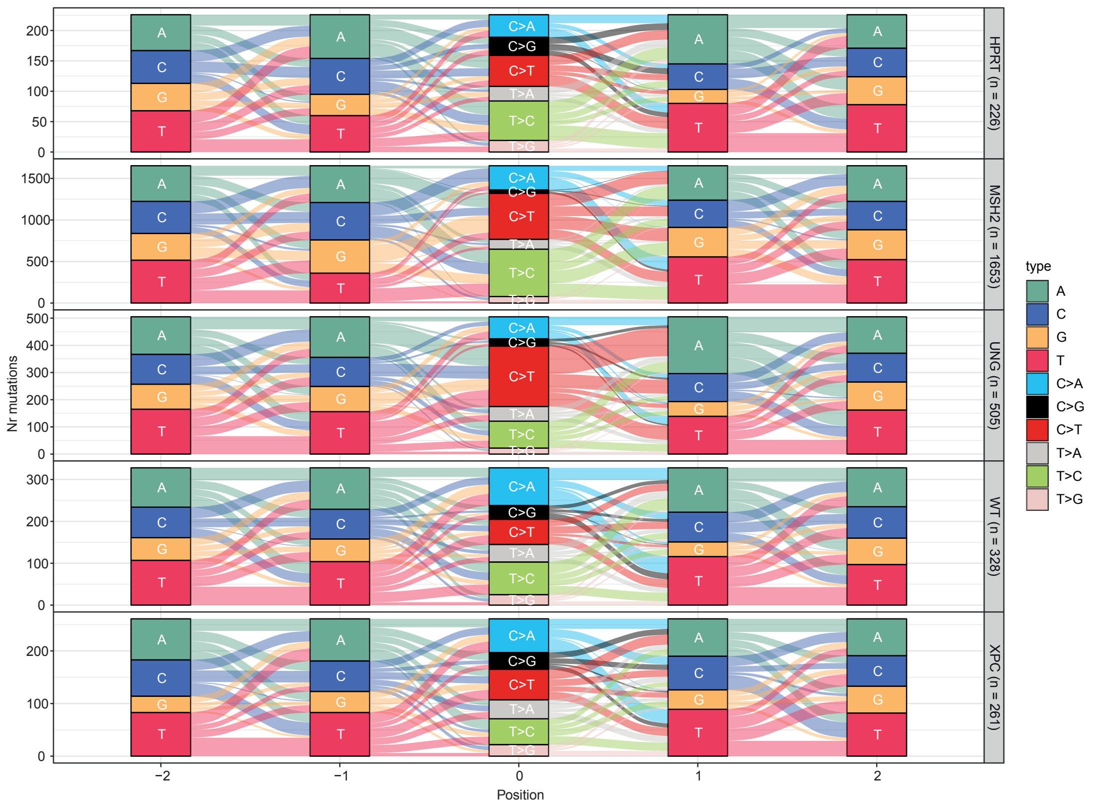


Fig. S2 Mutation contexts can be visualized with a river plot

A river plot depicting the indicated mutation types and the surrounding context for each sample. The bars show the number of mutations or bases for each type. The flows show the connections between the mutations and their context.

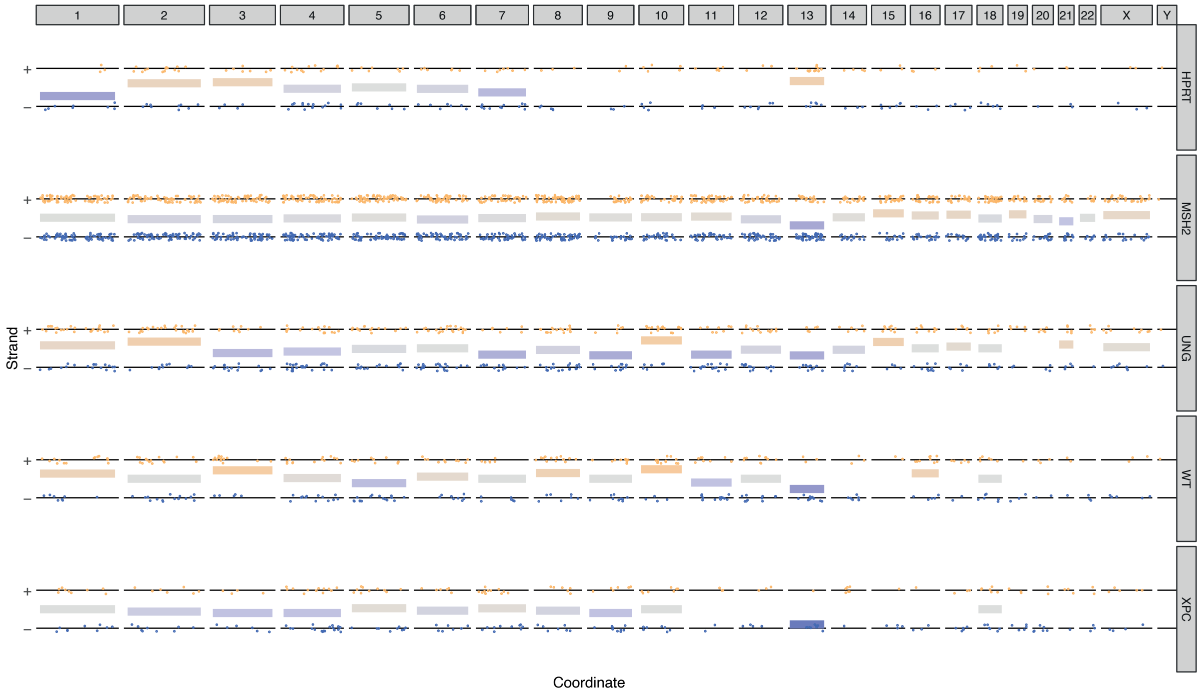


Fig. S3 Lesion segregation can be visualized

A jitter plot depicting the presence of lesion segregation for each sample per chromosome. Each dot depicts a single base substitution. Any C>N or T>N is shown as a “+” strand mutation, while G>N and A>N mutations are shown on the “-“ strand. The x-axis shows the position of the mutations. The horizontal lines are calculated as the mean of the "+" and "-" strand, where "+" equals 1 and "-" equals 0. They indicate per chromosome on which strand most of the mutations are located. In this example no lesion segregation was present.

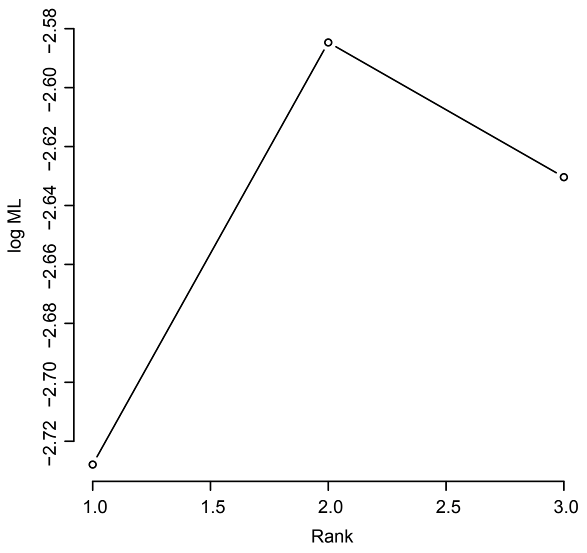


Fig. S4 Variational Bayes NMF can be used to predict the optimal number of signatures to extract

The log maximum likelihood is shown for different ranks. The rank is the number of signatures to extract. The highest likelihood shows the optimal rank. In this case this is 2.

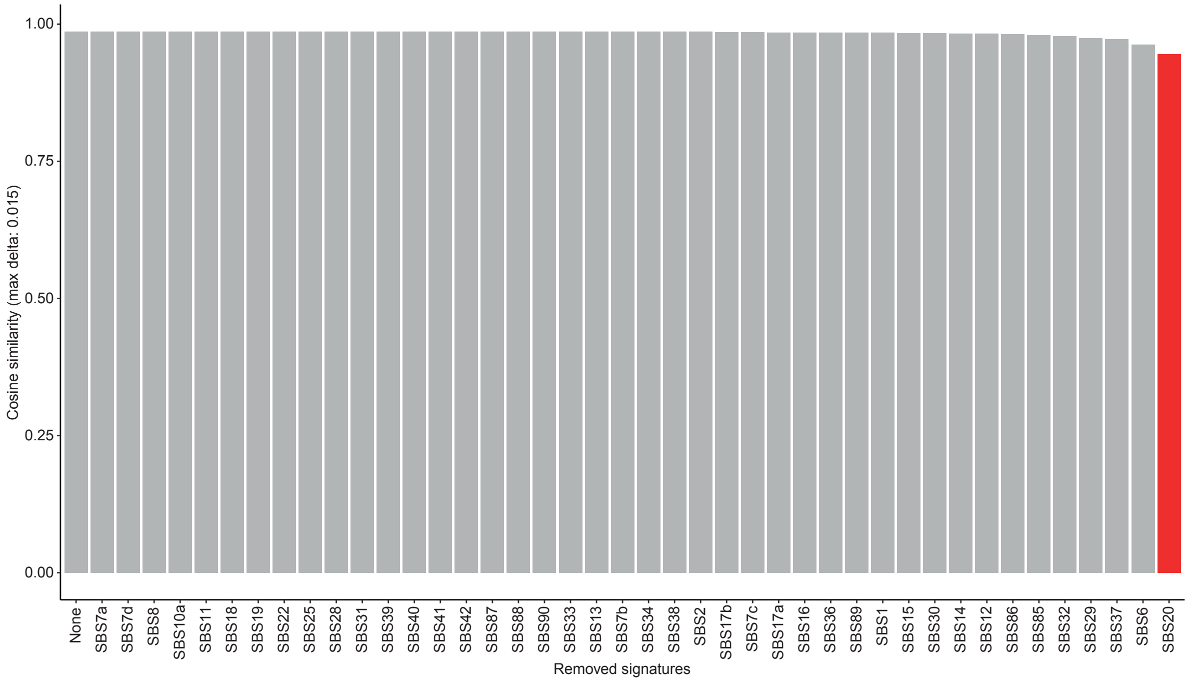


Fig. S5 Strict refitting iteratively removes signatures

The cosine similarity between the original and reconstructed profile of the *MSH2* knockout during different iterations of the strict refitting process. The signatures with the lowest contributions are iteratively removed and the cosine similarity is calculated. This is depicted from left to right. Removing SBS20 decreased the cosine similarity more than the cutoff, so it was retained, and the algorithm stopped.

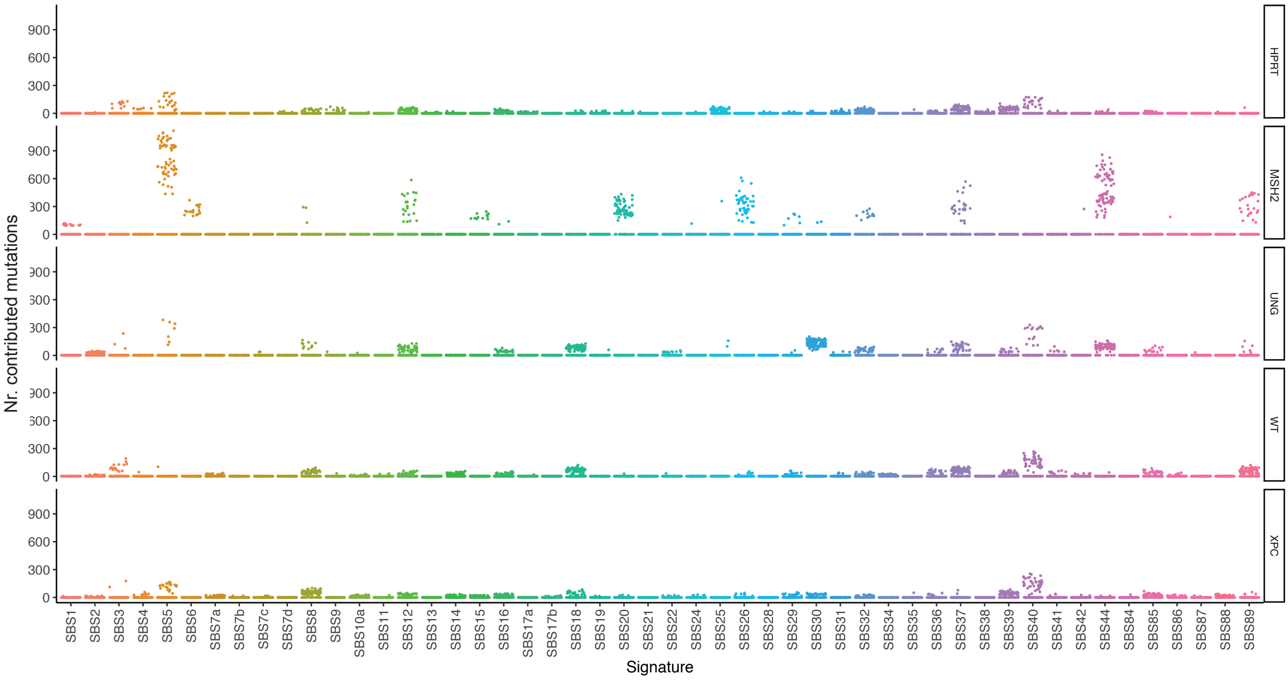


Fig. S6 Bootstrapped signature refitting can be visualized with a jitter plot

A jitter plot depicting the bootstrapped signature refitting for each sample. Each dot shows the number of mutations contributed by a signature according to one bootstrap iteration.

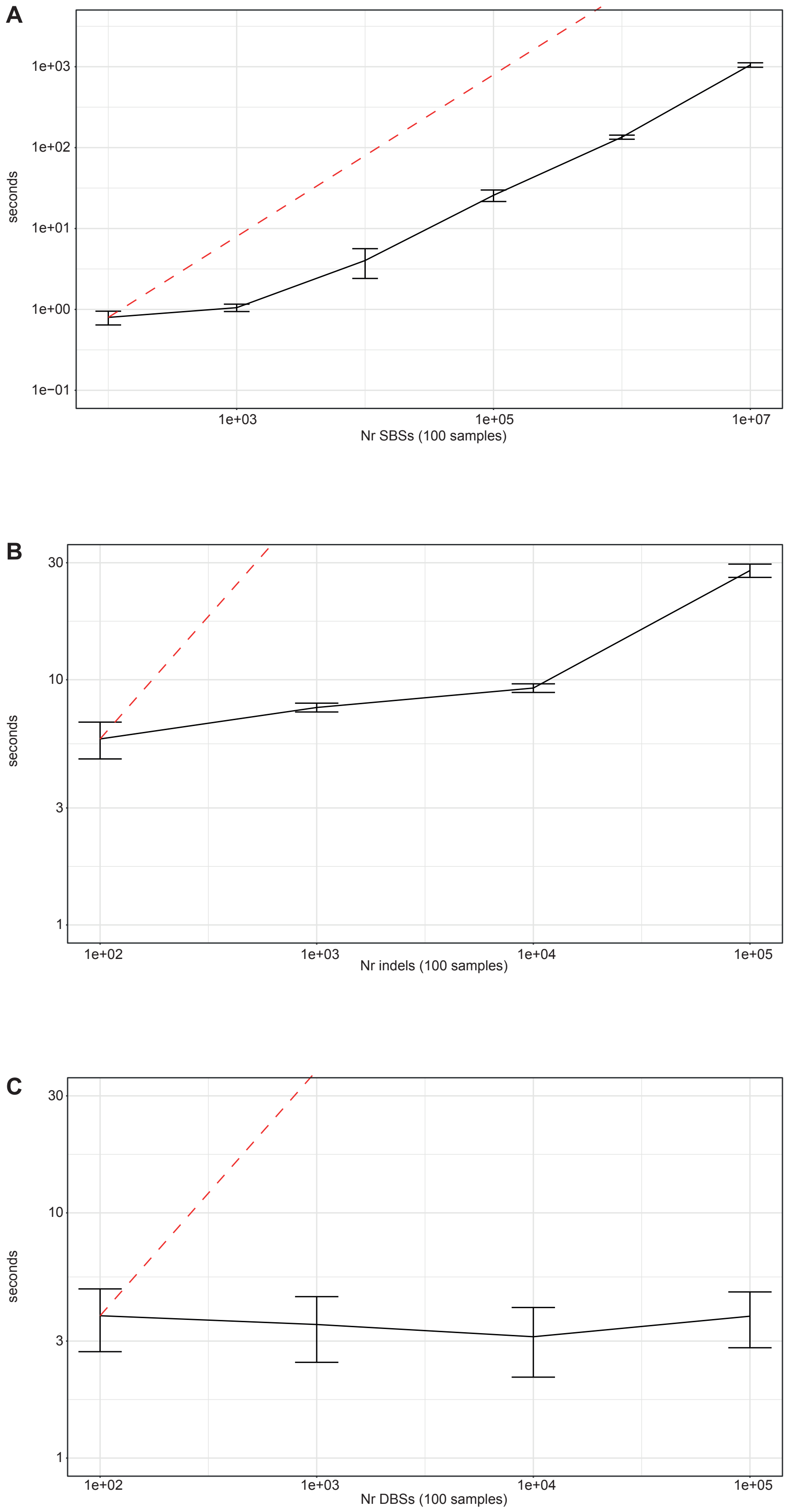


Fig. S7 The matrix generating functions have O(n) or better scaling

The y-axis shows the time it takes to generate a mutation matrix for the number of mutations on the x-axis for **a** SBSs, **b** indels and **c** DBSs. The mutations are always split over 100 samples. The dashed red line indicates O(n) scaling. The SBS and indel functions approach O(n) scaling on large mutations sets. The runtime of the DBS function is independent of the number of mutations.

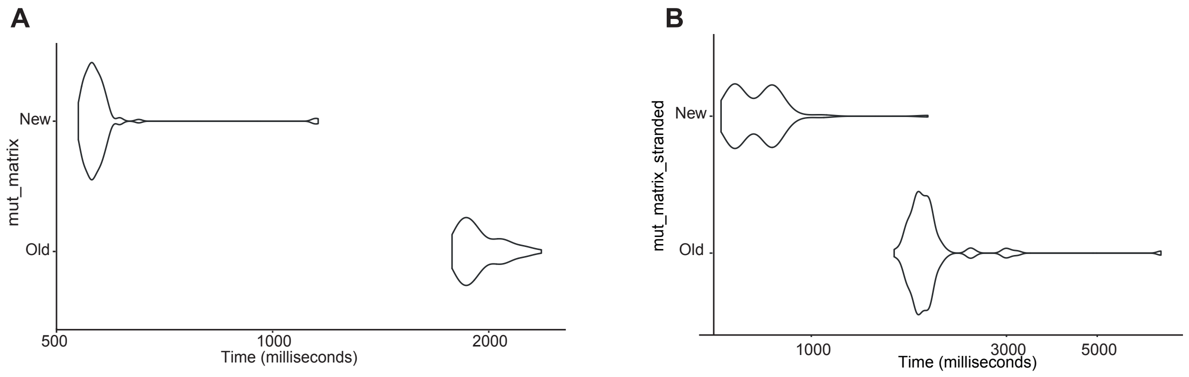


Fig. S8 Benchmark of the “mut_matrix” function

Violin plot depicting the run-times of the new and the old versions of the **a** “mut_matrix” and **b** “mut_matrix_stranded” functions. The benchmark was run on a 2019 MacBook Pro (2.4 GHz Quad-Core, 16GB RAM).

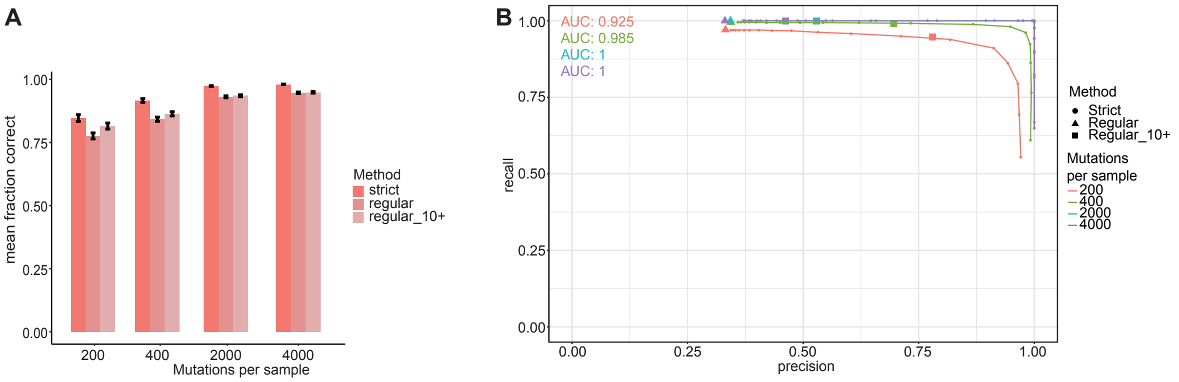


Fig. S9 Recall-precision plot of different refitting methods in MutationalPatterns

**a** Bar graph depicting the mean fraction of correctly attributed mutations for “regular”, “regular_10+” and “strict” refitting. This is shown for 4 experiments. Per experiment a mutation matrix with 300 simulated samples, each containing 4 signatures, was generated. The number of mutations per sample was respectively 200, 400, 2000 and 4000 for the 4 different experiments. The error bars show the 95% confidence interval. The fraction of correctly attributed mutations is calculated as 1 minus the absolute difference between the real and estimated contribution divided by the sum of the real and estimated contribution. **b** Recall-precision plot showing the recall (sensitivity) and precision of the “strict” method when different “max_delta” cutoffs are used for signature refitting. The recall and precision of the “regular” and “regular_10+” methods are also shown with respectively triangles and squares. Since these methods don’t have a “max_delta” cutoff only a single point can be shown for them. This is shown for 4 experiments. Per experiment a mutation matrix with 300 simulated samples, each containing 4 signatures, was generated. The number of mutations per sample was respectively 200, 400, 2000 and 4000 for the 4 different experiments. The area under the curve (AUC) is shown per experiment.

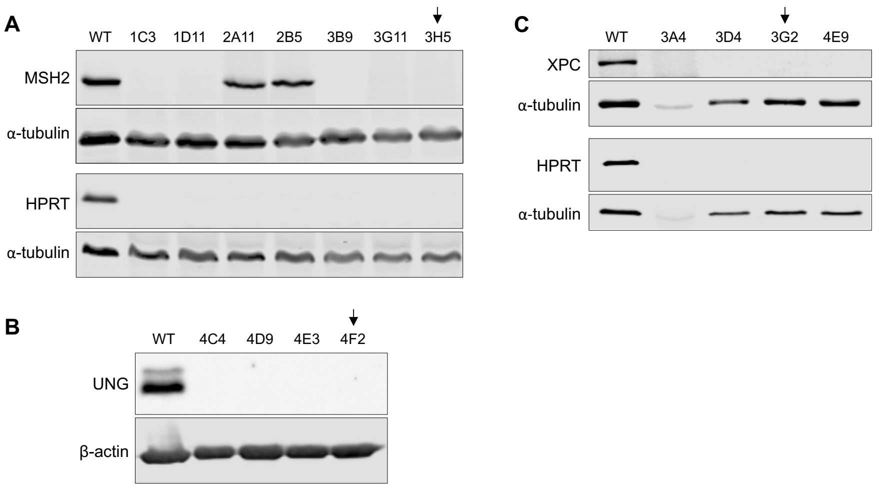


Fig. S10 Western blot analysis of AHH-1 CRISPR-Cas9 edited clonal lines

**a** Western blot of total protein lysate of bulk AHH-1 (WT) or single cell clones generated after transfection with CRISPR-Cas9 plasmids targeting *MSH2* and *HPRT*. All clones are HPRT knockout, and 5/7 clones are also knockout for MSH2. ***α***-Tubulin staining was done on the same membrane as the indicated protein above. **b** Western blot of total protein lysate of bulk AHH-1 (WT) or single cell clones generated after transfection with CRISPR-Cas9 plasmids targeting *UNG* and *HPRT*. All clones are knockout for UNG and HPRT (HPRT blot not shown). β-actin staining was done on the same membrane as UNG staining. **c** Western blot of total protein lysate of bulk AHH-1 (WT) or single cell clones generated after transfection with CRISPR-Cas9 plasmids targeting *XPC* and *HPRT*. All clones are HPRT and MSH2 knockout. ***α***-Tubulin staining was done on the same membrane as the indicated protein above. Arrows indicate clones selected for a second clonal step and whole genome sequencing.

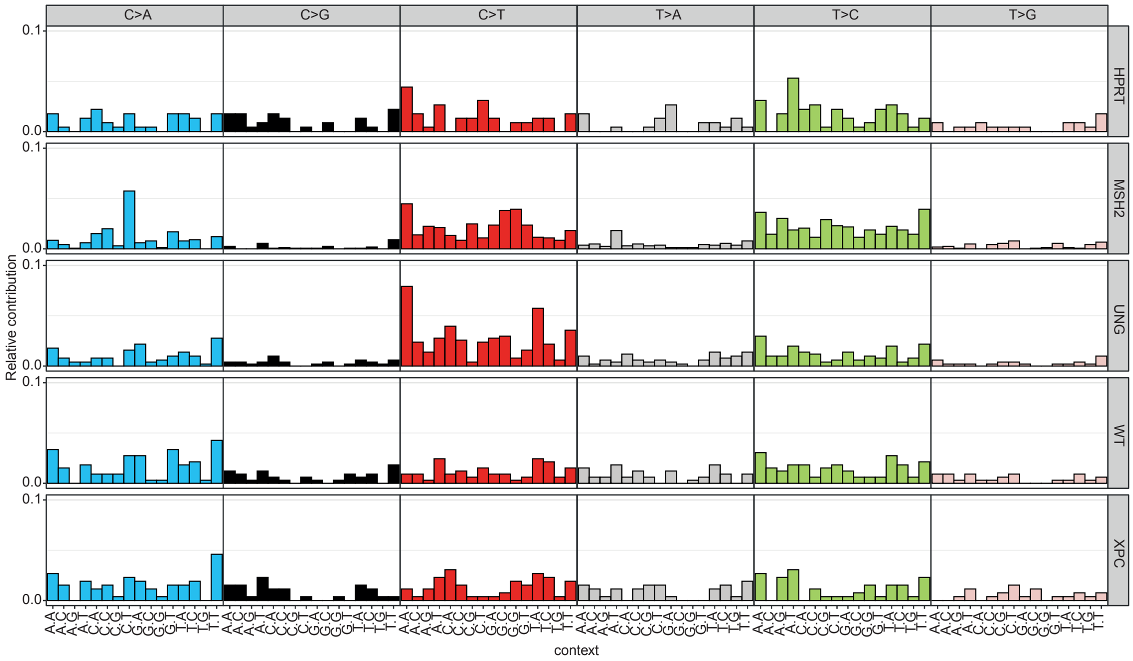


Fig. S11 SBS profiles of knockout samples

Relative contribution of each trinucleotide change to the point mutation spectrum for each sample.
