## Additional file 2 for "MutationalPatterns: The one stop shop for the analysis of mutational processes"

### Methods

#### Cell lines

AHH-1 cells, a human B lymphoblastoid cell line, were obtained from ATCC. Cells were cultured in RPMI 1640 medium (Gibco) with 2 mM L-glutamine, 1.5 g/L sodium bicarbonate, 4.5 g/L glucose, 10 mM HEPES, 1.0 mM sodium pyruvate, 10% heat-inactivated horse serum and penicillin/streptomycin. Low-passage cells were used for transfection experiments.

#### CRISPR-Cas9 gene editing

The human codon-optimized Cas9 expression plasmid was obtained from Addgene (px330-U6-Chimeric_BB-CBh-hSpCas9). The gRNA sequences were inserted by BbsI digestion and T4 ligation as described (Ran et al., 2013). sgRNA target sequences: hHPRT-sgRNA 5'-GGCTTATATCCAACACTTCG-3', hMSH2-sgRNA 5'-ACAAAGACTTGTTAACCAG-3', hUNG-sgRNA 5'-TCGGCACTCAGCGGCGAGGA-3', hXPC-sgRNA 5'-AAAGATTGACTGCGGATCC-3'. To generate DNA repair gene knockout lines, single cell suspensions of AHH-1 cells were co-transfected with Cas9- and sgRNA-expressing px330 plasmids, targeting *HPRT* and either *MSH2, UNG* or *XPC*. Plasmid DNA was mixed in an equal ratio and combined with Lipofectamine 2000. Transfection was performed according to manufacturer's instructions in AHH-1 medium with 1% horse serum. After 3 hours, complete AHH-1 medium (10% horse serum) was added to the cells and cells were cultured for 6 days. On day 6, 0.5 µg/mL 6-thioguanine (6-TG) was added to the cells after making single cell suspensions to select for *HPRT* knockout cells. On day 19, cells growing on 6-TG were plated in limiting dilutions to obtain single cell clones. Growing clones were analyzed by PCR and Western blot for knockout of *HPRT* and *MSH2, UNG* or *XPC*. Selected clones were subjected to another round of limiting dilutions to obtain subclones of the selected clones.

**Western blot**

Cell pellets were directly lysed in sample buffer (62.5 mM Tris-HCl, 2.5% SDS, 10% glycerol, 0.002% bromophenol blue, 100 mM DTT). Total protein lysates were loaded on SDS-PAGE gels and transferred to nitrocellulose membranes (BioRad). Membranes were blocked and probed with antibodies directed against HPRT (ab10479, Abcam), MSH2 (D24B5, Cell Signaling Technology), UNG (OTI1A11, ThermoFisher Scientific), XPC (12701, Cell Signaling Technology), β-actin (RM112, Sigma-Aldrich) and α-tubulin (B-5-1-2, Sigma-Aldrich).

**Whole genome sequencing and read alignment**

DNA libraries for Illumina sequencing were generated by using standard protocols (Illumina) from 500 ng of genomic DNA isolated from the clonally expanded AHH-1 cells using QIAamp DNA Blood & Tissue Kit (QIAGEN) according to manufacturers' instructions. All samples were sequenced (2 × 150 bp) by using Illumina HiSeq X Ten or NovaSeq 6000 sequencers to 30X base coverage. Whole genome sequencing data was mapped against human reference genome GRCh38 by using Burrows-Wheeler Aligner v0.7.5a mapping tool (Li and Durbin, 2010) with settings 'bwa mem -c 100 -M'. Sequence reads were marked for duplicates by using Sambamba v0.6.8 markdup. Full pipeline description and settings also available at: https://github.com/UMCUGenetics/IAP.

**Mutation calling and filtering**

Raw variants were multisample-called by using the GATK HaplotypeCaller v3.8-1-0 (Depristo et al., 2011) and GATK-Queue v3.8-1-0 with default settings and additional option

'EMIT_ALL_CONFIDENT_SITES'. The quality of variant and reference positions was evaluated by using GATK VariantFiltration v3.8-1-0 with options -snpFilterName SNP_LowQualityDepth -snpFilterExpression "QD < 2.0" -snpFilterName SNP_MappingQuality -snpFilterExpression "MQ < 40.0" -snpFilterName SNP_StrandBias -snpFilterExpression "FS > 60.0" -snpFilterName SNP_HaplotypeScoreHigh -snpFilterExpression "HaplotypeScore > 13.0" -snpFilterName SNP_MQRankSumLow -snpFilterExpression "MQRankSum < -12.5" -snpFilterName SNP_ReadPosRankSumLow -snpFilterExpression "ReadPosRankSum < -8.0" -snpFilterName SNP_HardToValidate -snpFilterExpression "MQ0 >= 4 && ((MQ0 / (1.0 * DP)) > 0.1)" -snpFilterName SNP_LowCoverage -snpFilterExpression "DP < 5" -snpFilterName SNP_VeryLowQual -snpFilterExpression "QUAL < 30" -snpFilterName SNP_LowQual -snpFilterExpression "QUAL >= 30.0 && QUAL < 50.0 " -snpFilterName SNP_SOR -snpFilterExpression "SOR > 4.0" -cluster 3 -window 10 -indelType INDEL -indelType MIXED -indelFilterName INDEL_LowQualityDepth -indelFilterExpression "QD < 2.0" -indelFilterName INDEL_StrandBias -indelFilterExpression "FS > 200.0" -indelFilterName INDEL_ReadPosRankSumLow -indelFilterExpression "ReadPosRankSum < -20.0" -indelFilterName INDEL_HardToValidate -indelFilterExpression "MQ0 >= 4 && ((MQ0 / (1.0 * DP)) > 0.1)" -indelFilterName INDEL_LowCoverage -indelFilterExpression "DP < 5" -indelFilterName INDEL_VeryLowQual -indelFilterExpression "QUAL < 30.0" -indelFilterName INDEL_LowQual -indelFilterExpression "QUAL >= 30.0 && QUAL < 50.0" -indelFilterName INDEL_SOR -indelFilterExpression "SOR > 10.0". To obtain high-quality somatic mutation catalogs, we applied postprocessing filters as described (Blokzijl et al., 2016). Briefly, we considered variants at autosomal chromosomes without any evidence from a paired control sample (the original bulk culture used to generate the mutant lines); passed by VariantFiltration with a GATK phred-scaled quality score R 100; a base coverage of at least

10X in the clonal and subclonal cultures, and paired control sample; mapping quality (MQ) of 60; no overlap with single nucleotide polymorphisms (SNPs) in the Single Nucleotide Polymorphism Database v146; and absence of the variant in a panel of unmatched normal human genomes (BED-file available upon request). We additionally filtered heterozygous base substitutions with a GATK genotype score (GQ) lower than 99 in clonal or paired control samples. A GQ score of 10 was used for homozygous variants. For indels, we filtered variants with a GQ score lower than 99 in both clonal or subclonal culture, or paired control sample. A GQ of 20 was used for homozygous reference variants. (Blokzijl et al., 2016; Jager et al., 2018). Finally, we only considered variants with a variant allele frequency of ≥0.3 in the sub-clones and a variant allele frequency lower than 0.3 in the original paired clones. These variants specifically accumulated between the two clonal expansion steps. The script is available at: https://github.com/ToolsVanBox/SMuRF.

**Additional references**

Blokzijl, F., de Ligt, J., Jager, M., Sasselli, V., Roerink, S., Sasaki, N., Huch, M., Boymans, S., Kuijk, E., Prins, P., et al. (2016). Tissue-specific mutation accumulation in human adult stem cells during life. Nature *538*, 260–264.

Depristo, M.A., Banks, E., Poplin, R., Garimella, K. V., Maguire, J.R., Hartl, C., Philippakis, A.A., Del Angel, G., Rivas, M.A., Hanna, M., et al. (2011). A framework for variation discovery and genotyping using next-generation DNA sequencing data. Nat. Genet. *43*, 491–501.

Jager, M., Blokzijl, F., Sasselli, V., Boymans, S., Janssen, R., Besselink, N., Clevers, H., van Boxtel, R., and Cuppen, E. (2018). Measuring mutation accumulation in single human adult stem cells by whole-genome sequencing of organoid cultures. Nat. Protoc. *13*, 59–78.

Li, H., and Durbin, R. (2010). Fast and accurate long-read alignment with Burrows-Wheeler transform. Bioinformatics *26*, 589–595.

Ran, F.A., Hsu, P.D., Wright, J., Agarwala, V., Scott, D.A., and Zhang, F. (2013). Genome engineering using the CRISPR-Cas9 system. Nat. Protoc. *8*, 2281–2308.
